# The evolutionary dynamics of microRNAs in domestic mammals

**DOI:** 10.1101/257006

**Authors:** Luca Penso-Dolfin, Simon Moxon, Wilfried Haerty, Federica Di Palma

## Abstract

MicroRNAs are crucial regulators of gene expression found across both the plant and animal kingdoms. While the numberof annotated microRNAs deposited in miRBase has greatly increased in recent years, few studies provided comparative analyses across sets of related species, or investigated the role of microRNAs in the evolution of gene regulation.

We generated small RNA libraries across 5 mammalian species (cow, dog, horse, pig and rabbit) from 4 different tissues (brain, heart, kidney and testis). We identified 1675 miRBase and 413 novel microRNAs by manually curating the set of computational predictions obtained from *miRCat* and *miRDeep2*.

Our dataset spanning five species has enabled us to investigate the molecular mechanisms and selective pressures driving the evolution of microRNAs in mammals. We highlight the important contributions of intronic sequences (366 orthogroups), duplication events (135 orthogroups) and repetitive elements (37 orthogroups) in the emergence of new microRNA loci.

We use this framework to estimate the patterns of gains and losses across the phylogeny, and observe high levels of microRNA turnover. Additionally, the identification of lineage-specific losses enables the characterisation of the selective constraints acting on the associated target sites.

Compared to the miRBase subset, novel microRNAs tend to be more tissue specific. 20 percent of novel orthogroups are restricted to the brain, and their target repertoires appear to be enriched for neuron activity and differentiation processes. These findings may reflect an important role for young microRNAs in the evolution of brain expression plasticity.

Many seed sequences appear to be specific to either the cow or the dog. Analyses on the associated targets highlight the presence of several genes under artificial positive selection, suggesting an involvement of these microRNAs in the domestication process.

Altogether, we provide an overview on the evolutionary mechanisms responsible for microRNA turnover in 5 domestic species, and their possible contribution to the evolution of gene regulation.

## Introduction

MicroRNAs (miRNAs) are short, ~22 nt non-coding RNA molecules found across the plant and animal kingdoms. They represent important regulators of gene expression which have been shown to be implicated in fundamental processes such as embryonic development or tissue differentiation (Lee et al. 1993;)(Brennecke et al. 2003;)(Xu et al. 2003;)(Bushati et al. 2008). microRNAs typically act by binding to complementary RNA molecules, resulting in translational repression or mRNA degradation (Bartel 2004;) (Niwa and Slack 2007); (Carthew and Sontheimer 2009). Their biogenesis starts with the transcription of a long RNA molecule, the *pri-miRNA*, located inside the nucleus of a cell. This precursor, characterized by one or more stem-loop structures, is processed by the enzyme *Drosha* which cleaves the double-stranded stem region. The resulting *pre-miRNA* is then exported out of the nucleus; following the excision of the loop region operated by *Dicer*, a ~22 bp, double stranded RNA molecule will be generated (Berezikov 2011;) Ha and Kim 2014). One of these two strands (referred as 5p-and 3p-miRNA) will be typically degraded (Ha and Kim 2014), while the other will be loaded into the miRNA-induced silencing complex and guide the targeting of mRNA molecules, by partial base-pairing (Berezikov 2011).

The recent advent of *RNA-Seq* technology (Wang et al. 2009) and the increasing number of assembled genomes provide us with greater power to study microRNA function and evolution. Computational tools based on this technology have been recently developed, allowing for an *in silico* identification of putative microRNA loci from a genome assembly and small RNA reads data for the same species (Friedlander et al. 2012); (Stocks et al. 2012;)(Wu et al. 2013;)(Zheng et al. 2016;)(Paicu et al. 2017). Studies based on homology analyses and computational microRNA prediction have allowed for the recent identification of thousands of microRNAs, available online from databases such as miRBase (Griffiths-Jones 2006).

While many studies have been focusing on the functional role of microRNAs, especially in disease, few have tried to clarify their evolutionary history. As microRNAs represent a relatively easy path to phenotypic diversification, through both temporal and tissue specific variations in gene expression, there is a great interest in elucidating their evolution including gains and losses, and how in turn this relates to gene regulation and target sites evolution.

Meunier *et al.* (Meunier et al. 2013) highlighted the high rates of miRNA family gains in placentals and marsupials, and the key role of introns and duplication events in the emergence of novel miRNA loci. Their analyses also suggested a gradual increase in expression levels for selectively retained microRNA families, along with changes in target repertoires, while many novel microRNAs with neutral or deleterious regulatory effects seem to be rapidly lost. Flynt *et al.* (Alex Flynt 2017) provided an overview of the microRNA diversity and evolution in the *Drosophila* genus. The authors generated a new microRNA annotation across 11 species, supported by deep sequencing from multiple tissues. They inferred gain and loss patterns across the *Drosophila* phylogeny, described cases of clade specific, 5’ end shift in miRNA processing, and compared different subpopulations of their large set of novel microRNA loci.

Other studies focused on the evolution of 3’UTR target sites, looking at their conservation across different evolutionary timescales. Xu *et al* (Xu et al. 2013) used high confidence CLIP data to define the evolvability of microRNA targets in vertebrates. They found that the conservation levels progressively decrease as larger taxonomic groups are considered, with 94% of target sites being conserved among Human and Chimpanzee, 80% among Human and 10 other Mammalian species, and only 6% between Human and Zebrafish. Chen and Rajewsky ((Chen and Rajewsky 2006) observed small numbers of conserved target sites across vertebrates, flies and nematodes. However, they were able to identify a small subset of deeply conserved target sites, and pointed out the enrichment for developmental processes in the corresponding genes. Comparative analyses performed by Friedman et al *(Friedman et al. 2009)*, on the contrary, suggest that a high number of predicted 3’ UTR target sites are conserved above background levels in mammals. However, results might have been influenced by the use of sequence conservation (Pct score) as one of the criteria for *in silico* target identification.

In this study, we focus on the evolution of microRNAs in five domestic species of great economic and biomedical interest: cow, dog, horse, pig and rabbit, none of which have been previously included in a comparative study across domestic mammals.

The cow (*Bos taurus*) and the pig (*Sus scrofa*) represent invaluable resources for food production (Russo et al. 2002;) (Bovine Genome et al. 2009;) (Bendixen et al. 2010;) (Seo et al. 2013;) Schook et al. 2015). There is a great economical interest in gaining more understanding about the genetic basis of agro-economically relevant traits (for example, milk productivity, resistance to pathogens, stress, meat quality) (Russo et al. 2002). Moreover, the pig’s high resemblance to humans in anatomy, physiology and genetics has also encouraged recent biomedical research (Bendixen et al. 2010;)(Groenen et al. 2012;)(Walters et al. 2012;)(Schook et al. 2015).

The dog (*Canis familiaris*) is a model system for several human diseases, and a unique example of great phenotypic diversification following a domestication event. Abundant polymorphism data have been generated (Lindblad-Toh et al. 2005), while GWAS studies on this organism have successfully identified the genetic base of heritable diseases (Awano et al. 2009); (Lei et al. 2009;)(Wilbe et al. 2010;)(Tonjes et al. 2014;) (Bartolome et al. 2015;) (Brinkmeyer-Langford et al. 2016;)(Truve et al. 2016). Genetic and genomic studies on the horse (*Equus caballus*) have mainly aimed at understanding the biology of infectious, respiratory and allergic diseases these animals are subject to, and the development of adequate therapies (https://www.uky.edu/Ag/Horsemap/welcome.html). However, the similarity with the corresponding human diseases means these studies have an even broader range of potential applications.

The rabbit (*Oryctolagus cuniculus*) represent yet another important model organism for biomedical studies, which has been used in research fields such as embryology, toxicology, pulmonary and cardiovascular research, as well as neurology (Craig et al. 2012).

To the best of our knowledge, this is the first comprehensive, comparative analysis of microRNA and target evolution across five domestic mammals, enabling us to investigate their potential role in the process or aftermath of domestication. We generate an improved microRNA annotation in these species, supported by deep sequencing from four different tissues (brain, heart, kidney and testis) and use this data to elucidate: 1) the relative contributions of different evolutionary mechanisms by which microRNAs newly arise; 2) the patterns of expression and gain/loss evolution of microRNA orthogroups, and their variation across different microRNA subpopulations; 3) the association of microRNA evolution with the regulation of specific biological processes, and their potential involvement in domestication; 4) the effect of branch specific microRNA loss on the conservation of the associated target sites; 5) the levels of target site conservation compared to the surrounding 3’ UTR regions.

## Results

### Improved genomic annotations of conserved and novel microRNA loci across five mammals

We ran miRCat (Stocks et al. 2012) and *miRDeep2* (Friedlander et al. 2012) on the corresponding combined set of small RNA libraries, thus generating two independent sets of putative miRNA loci. Predictions were then filtered based on the following two main criteria: evidence of both miRNA-3p and miRNA-5pexpression in small RNA reads alignments against the predicted loci, and miRNA - like hairpin secondary structure (as predicted using the *Vienna-RNA* package)(Hofacker 2009). This lead to the identification of a final set (union of miRCat and miRDeep2 high confidence predictions) of 2088 loci: 1675 miRBase annotated and 413 representing novel microRNA loci (Table 1). Tissue specific expression plots for all novel and conserved miRNAs were also generated (figg. S1-S5). The number of novel microRNA loci is particularly high in the dog. The computational predictions are highly dependent on sequencing depth, and in our study more brain samples were available for the dog compared to all the other species. Therefore, we sampled 40,680,089 reads (the total number of reads available for the horse) from the combined set of dog reads (n=99,597,072), and recalled dog microRNA loci. Based on our filtering criteria (coverage of at least one read at both the 3’ and 5’ ends, and a total coverage of at least 10 reads), we could still confidently annotate 111 novel loci. This corresponds to 66% of the original novel annotation, and still represents the highest count across all our species.

**Table 1.**
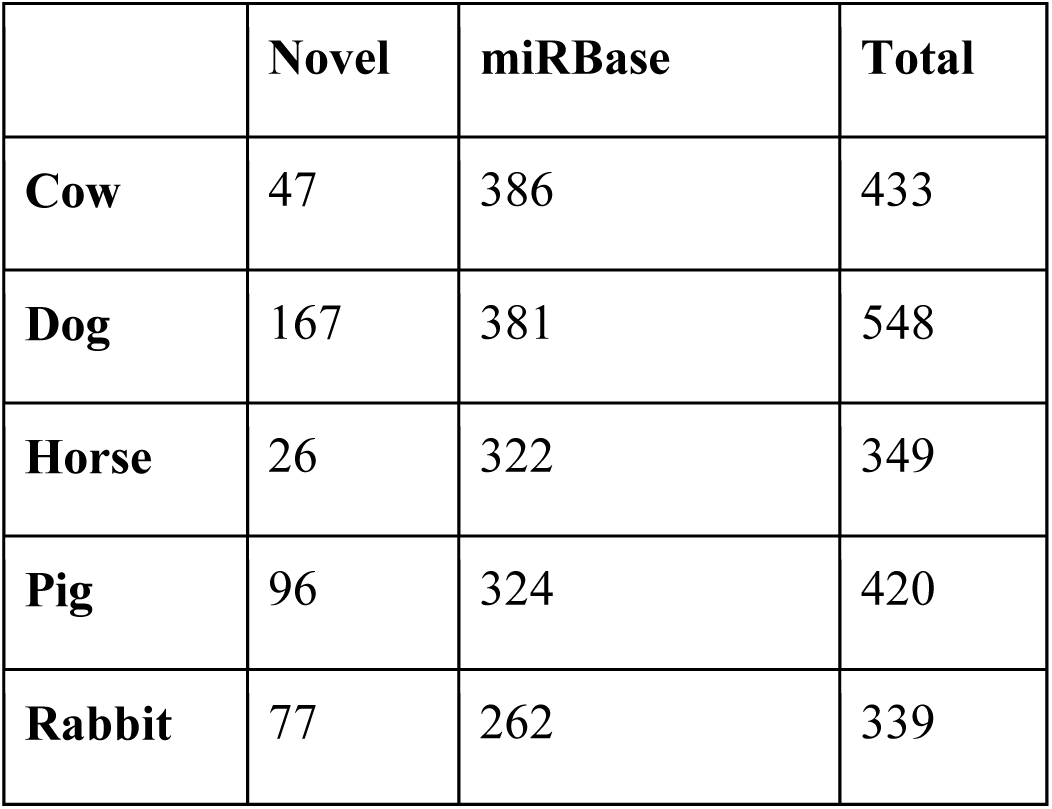
Counts of annotated microRNA loci, either belonging to a miRBase family or representing a novel gene.

We compared each of the five microRNA annotations with the corresponding latest *Ensembl* gene annotation (*B. taurus* UMD3.1.8, *C. familiaris* 3.1.86, *E. caballus* 2.86, *S. scrofa* 10.2.84, *O. cuniculus* 2.0.84). Unsurprisingly, we observed very high proportions of intronic and intergenic microRNAs, and very low numbers of microRNA overlapping UTR sequences. These patterns appear to be consistent across all 5 species (Fig.1). Analyses performed with *GAT* (Heger et al. 2013) indicate a significantly higher than expected overlap with introns in all species (q-value<0.001), while intergenic regions, despite containing a high number of microRNAs, are significantly underrepresented in all five genomes (q-value<0.001).

**Fig.1.**
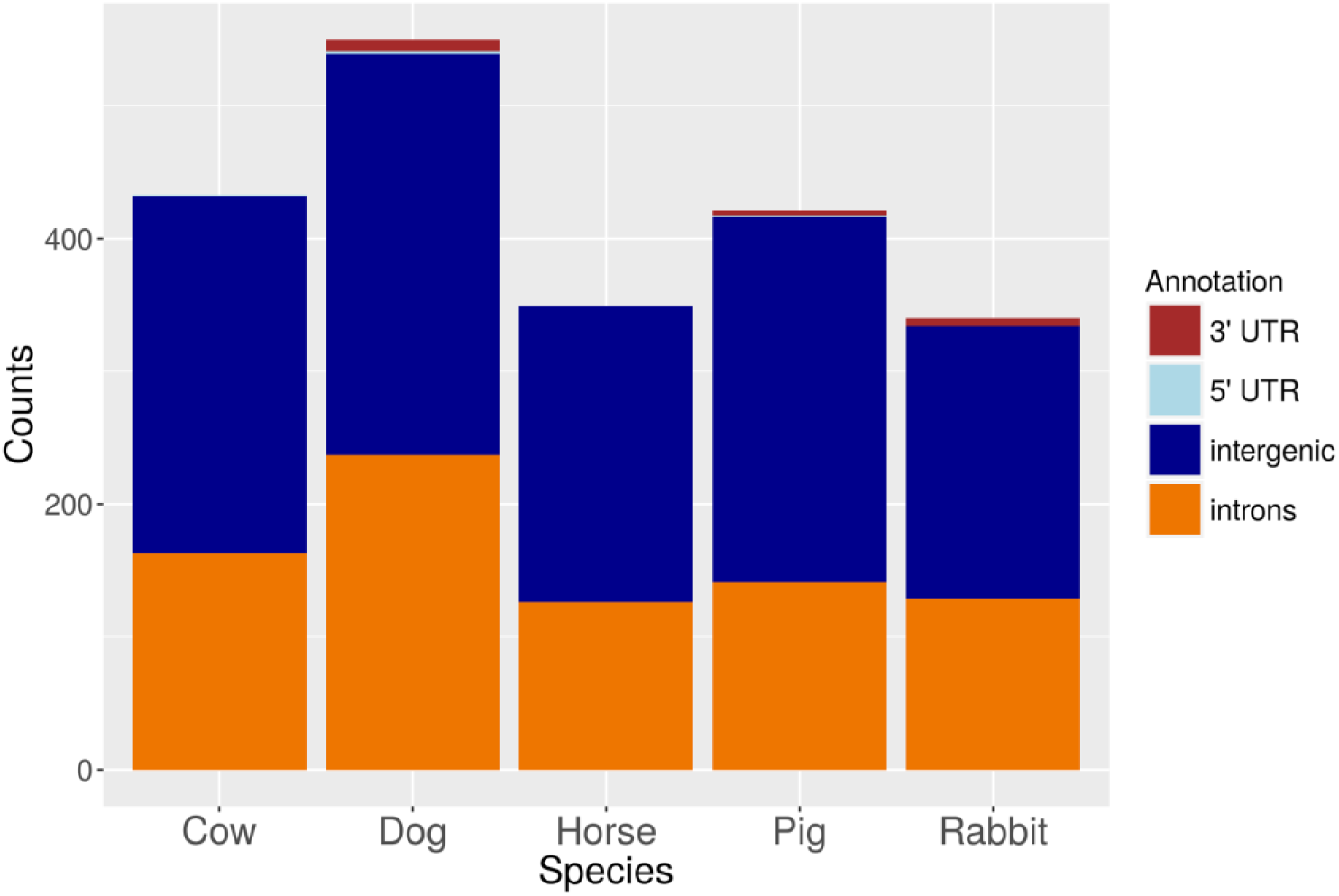
Genomic annotation for all predicted microRNA loci

### The evolution of mammalian microRNA families

In order to generate clusters of homologous microRNA loci, we used *CDhit* (Fu et al. 2012) with 80% minimum identity on our set of annotated microRNAs. We thus obtained a total of 732 clusters (orthogroups), representing 441 miRBase annotated and 291 novel (not present in miRBase) orthogroups.

In order to limit potential biases in our annotations resulting from different genome assembly quality and sequencing depth across our species, we decided to look for evidence of sequence homology between genome assemblies. We thus aligned all annotated miRNA loci against the genome assemblies of human, mouse and all species considered in this study. This allowed for the identification of loci missing in the annotation, but showing high sequence homology to a microRNA annotated in another species, as well as evidence of synteny conservation in the surrounding region (see Materials and Methods). As an additional strategy to overcome differences in annotation and assembly quality, we looked for annotated miRNA sequences in the set of unaligned reads of horse and pig (for which the initial estimates of microRNA gain rates were surprisingly low). This analysis allowed us to further improve the presence-absence information used for the gain/loss inference. We used this curated set to characterise the most parsimonious patterns of gain and loss of microRNA orthogroups across the phylogeny. Despite the high proportion of broadly conserved microRNA families, we observe high levels of microRNA turnover across the phylogeny, with many families being gained or lost in internal and terminal branches (fig. 2). We observe a positive net gain rate (dog, cow and rabbit) in all terminal branches, with the only exception being represented by the horse. The virtually equal rate of gain and loss is likely due to a lack of sequencing depth, resulting in a limited number of predicted novel, horse specific microRNAs.

**Fig 2.**
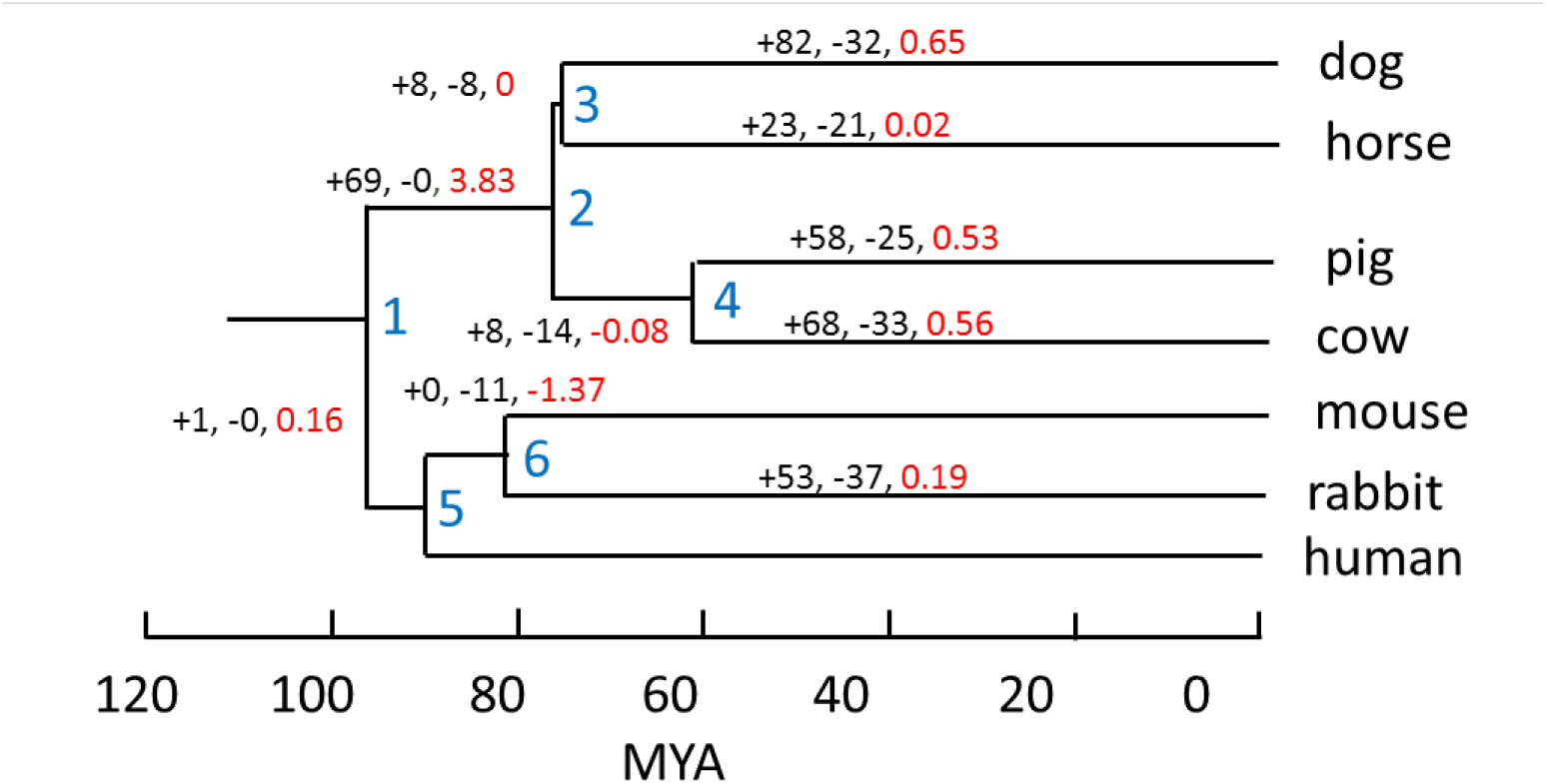
Gain and loss of microRNA clusters across the phylogenetic tree, as inferred by Dollo parsimony and synteny analyses

### Genomic sources of microRNAs

Various mechanisms can contribute to the appearance of new microRNA genes. While introns represent a crucial source of newly processed microRNA hairpins (sometimes not requiring Drosha processing, see the case of miRtrons), other microRNAs could arise by gene duplication, transcription of the opposite strand of an existing microRNA locus, or evolution from repetitive elements (Berezikov 2011). Our dataset allows us to determine the extent of the contribution of different evolutionary processes in mammals.

For the identification of microRNAs derived from repetitive elements, we used BLASTN (Altschul et al. 1990) to align all hairpin sequences against the *Repbase* (www.girinst.org/repbase) database. A bit score threshold, determined through the alignment of microRNAs against shuffled Repbase sequences (see Materials and Methods) was used in combination with other parameters to select high confidence BLAST hits. Following these conservative approach, we identified 73 novel and 45 miRBase microRNA loci showing a significant similarity with one or more Repbase sequences. Interestingly, 49 out of the 73 novel, putatively repeat-derived microRNA loci are part of a single, large orthogroup specific to the dog: cluster 508. We performed GO:term enrichment analyses on repeat-derived, dog specific microRNAs, and found significant enrichment for immunological processes (including “positive regulation of memory T cell differentiation”, GO:0043382; “positive regulation of activated T cell proliferation”, GO:0042104), as well as cognitive and behavioural (including “cognition”, GO:0050890; “behaviour”, GO:0007610, “exploration behaviour”, GO:0035640). These results might reflect an important role of these novel microRNAs in the *Carnivora* evolution of immune response and neural expression plasticity.

While we find 16 broadly conserved orthogroups, there are also 18 orthogroups appearing in terminal branches (Fig 3), suggesting an important role for repetitive elements in the emergence of novel microRNA loci.

**Fig 3.**
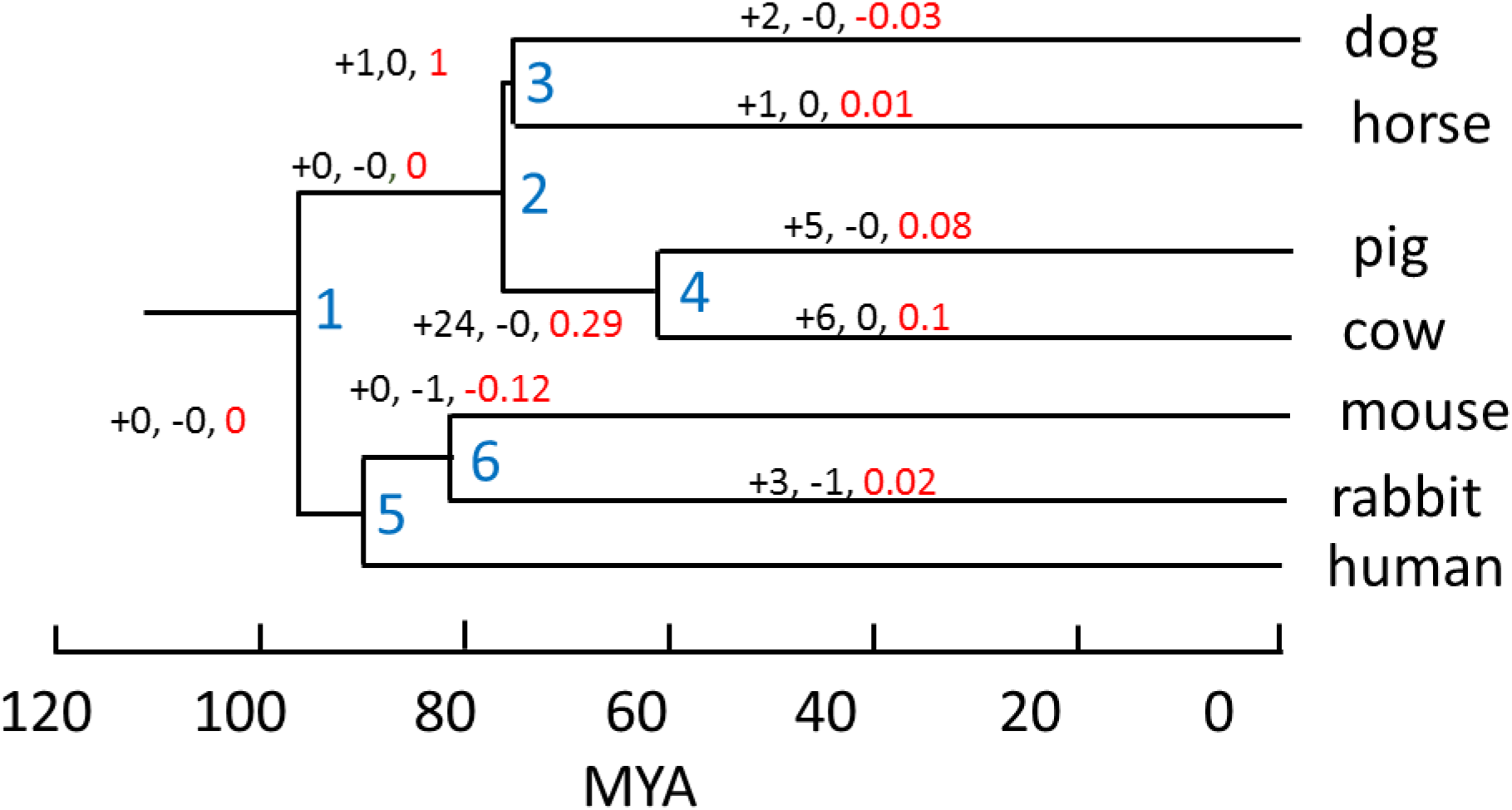
Gain and loss of repeat-derived microRNA clusters across the phylogenetic tree, as inferred by Dollo parsimony.

Next, we asked the question whether we could find any case of reverse complement microRNA sequences, lying on the opposite strand of exactly the same genomic interval. Our analyses lead to the identification of at least one of such cases in every species, with numbers ranging from 1 (in rabbit) to 22 coupled loci (in dog).

We also investigated microRNA clusters containing multiple paralogous gene copies, and identified a total of 135 orthogroups associated with duplication events (58 in tandem) in at least one species. We identified both old duplication events ( many duplicated loci have orthologous counterparts in other species) as well as more recent duplications, with genes arising in terminal branches (fig 4). Thus, duplication seems to be an important mechanism for the evolution of microRNAs in our set of species, as previously suggested by other studies in mammals (Meunier et al. 2013). When we compared the set of clusters with duplicated loci and with high similarity to repetitive elements, we identified 10 common clusters, 5 of which represents species specific groups.

**Fig 4.**
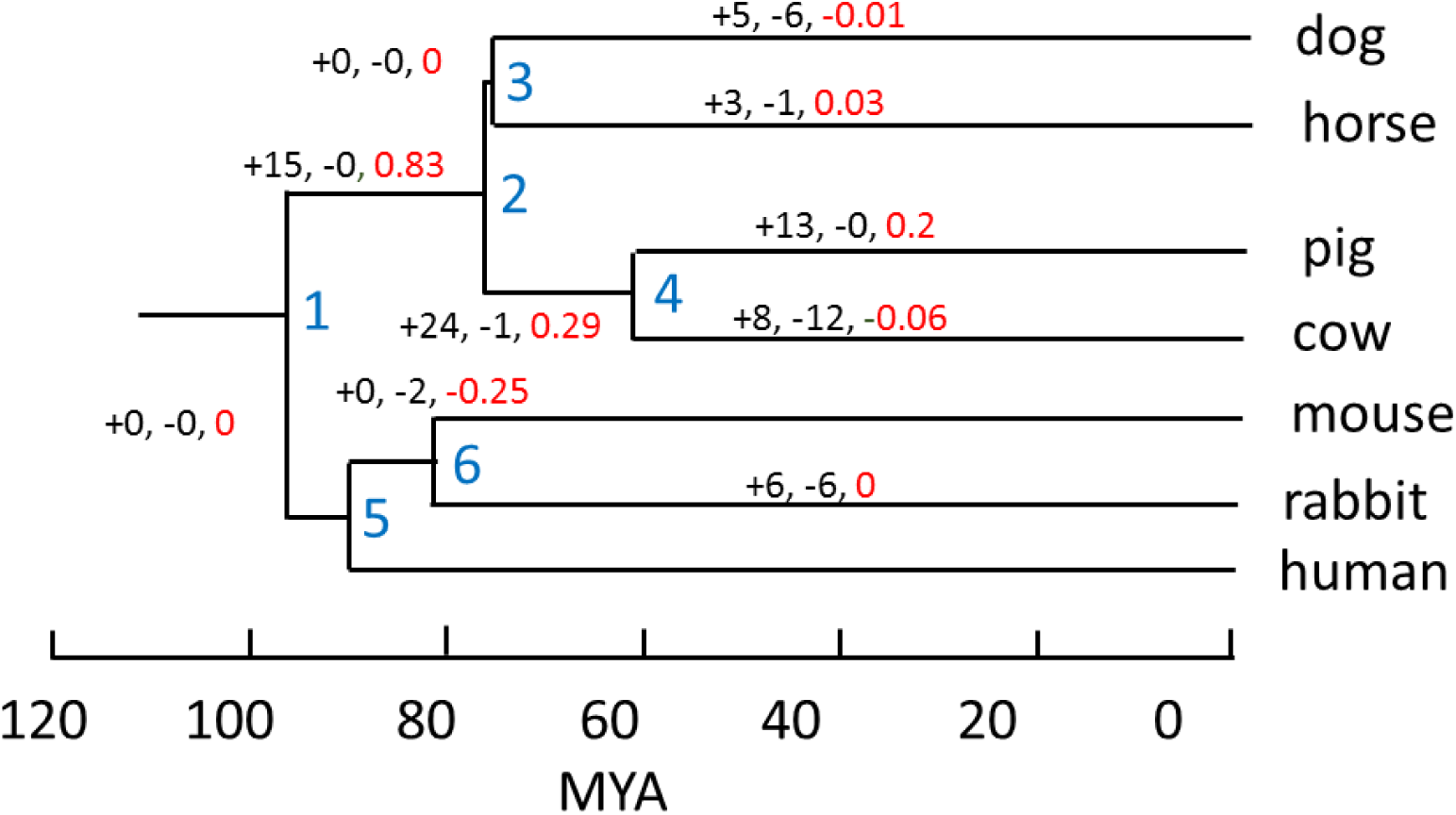
Gain and loss of clusters of duplicated microRNAs across the phylogenetic tree.

**Table 2.**
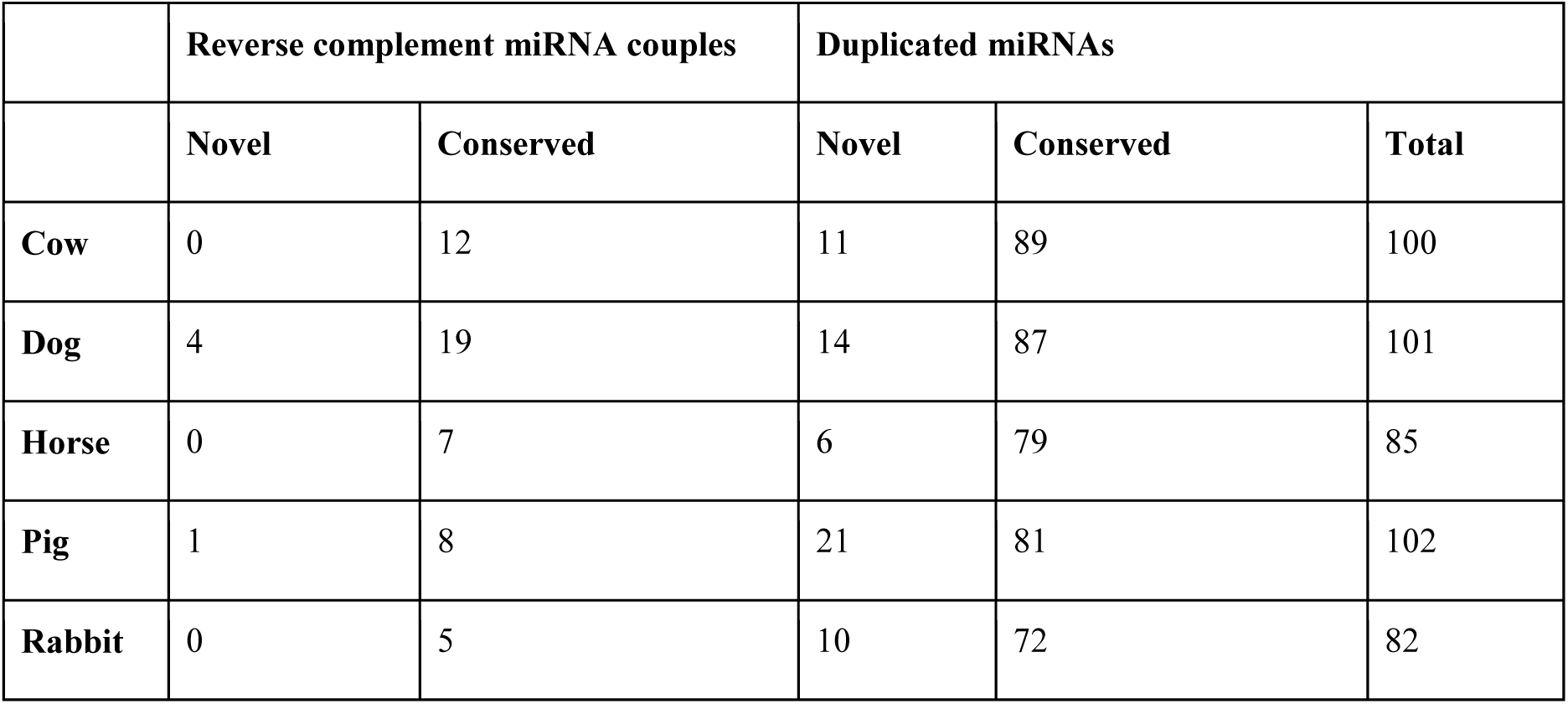
The number of reverse complement microRNA gene couples and number of miRNA orthogroups containing
paralogous duplicated genes

**Table 3.**
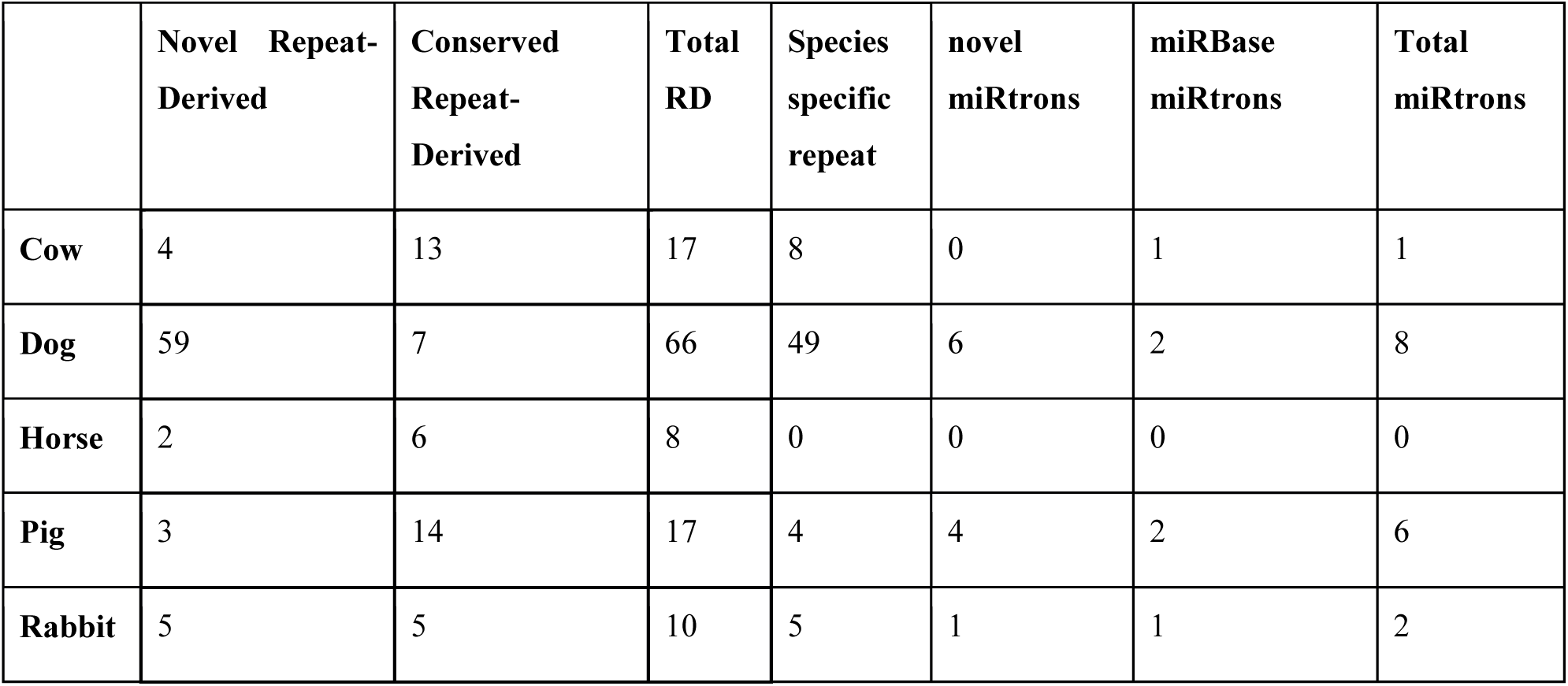
The number of miRtrons and putatively repeat-derived microRNA genes.

|  | Novel Repeat-Derived | Conserved Repeat-Derived | Total RD | Species specific repeat | novel miRtrons | miRBase miRtrons | Total miRtrons |
| --- | --- | --- | --- | --- | --- | --- | --- |
| **Cow** | 4 | 13 | 17 | 8 | 0 | 1 | 1 |
| **Dog** | 59 | 7 | 66 | 49 | 6 | 2 | 8 |
| **Horse** | 2 | 6 | 8 | 0 | 0 | 0 | 0 |
| **Pig** | 3 | 14 | 17 | 4 | 4 | 2 | 6 |
| **Rabbit** | 5 | 5 | 10 | 5 | 1 | 1 | 2 |

### The evolution of microRNA expression

Our dataset provides us with a great opportunity to investigate how the gain and loss of microRNAs relates to the observed patterns of miRNA expression across 4 different tissues. Our data suggests that young orthogroups tend to be more tissue restricted than the older, conserved ones, particularly when we consider brain and testis. (Figure 5) shows the number of gains across the phylogeny when considering only the orthogroups with evidence of expression in a particular tissue. By constructing trees where branch lengths are proportional to the fraction of brain specific orthogroups, we can clearly visualise the higher tissue restriction of young microRNAs. This observation is in line with previous findings, showing high proportions of novel microRNAs expressed in brain tissues (Meunier et al. 2013).

**Fig 5.**
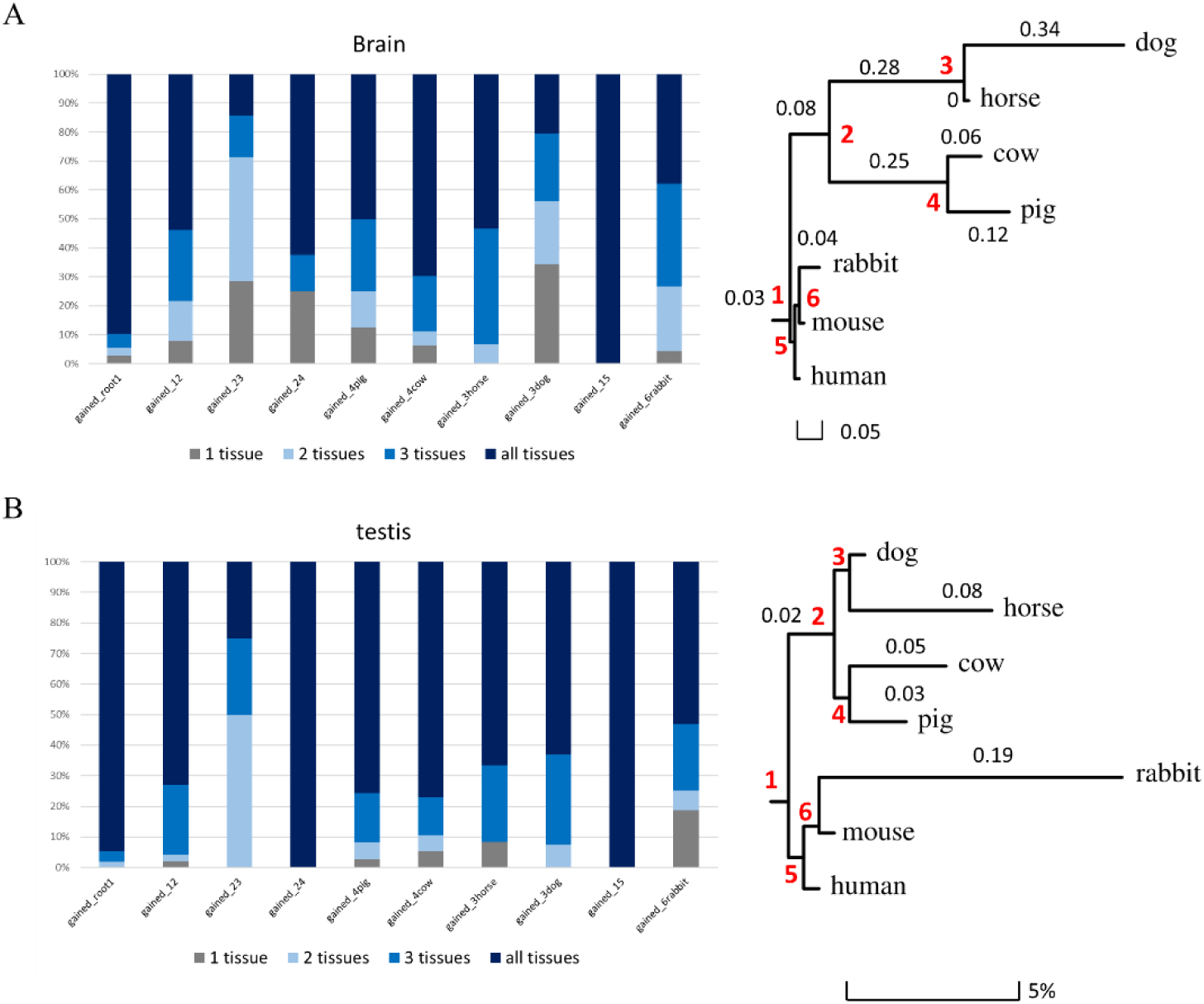
Expression patterns across the phylogeny for all microRNA orthogroups expressed in brain (A) and testis (B). Colour labelling indicate the number of microRNA families expressed in a specific tissue, divided in the following categories: tissue specific; expressed in the tissue + *n* additional ones (n=3 indicated as “all tissues”). Branch lengths in the phylogenetic trees are proportional to the fraction of tissue specific orthogroups.

Next, we asked the question whether we can see differences in tissue specificity between novel and miRBase microRNAs.Figure 6 shows the difference in the proportions of orthogroups expressed in a particular tissue, divided into sub-sets depending on the total number of tissues showing evidence of expression. We observed a significantly higher proportion of tissue specific families in the novel set compared to the miRBase set (z test, 03B1;=0.05) for all four tissues except the kidney, with a particularly striking difference in the cases of brain and testis (brain: p<10e^−4^, heart: p=0.012, kidney: p=0.051, testis: p=10e^−3^).

**Fig.6.**
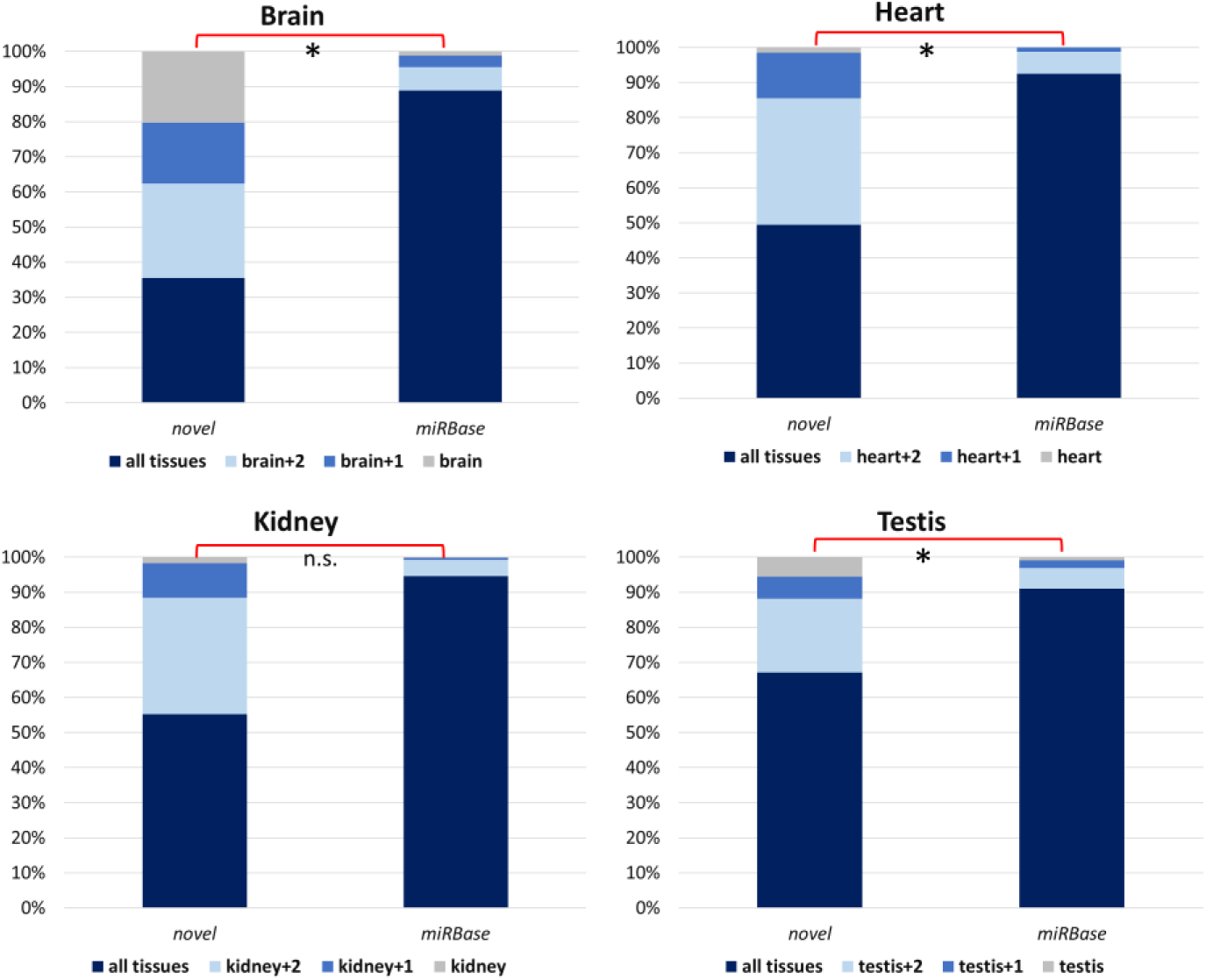
Expression patterns of novel and miRBase loci across 4 tissues. Colour labelling indicate the number of microRNA
families expressed in a specific tissue, divided in the following categories: not expressed in the tissue considered; tissue specific; expressed in the tissue + n additional ones (n=3 indicated as “all tissues”).

This result suggests that young microRNAs are expressed in a single or a few tissues when they first appear, and become more broadly expressed over time. Interestingly, we find that as much as 20% of novel orthogroups are restricted to the brain tissue. When we performed GO term enrichment analyses of the mRNA targets of novel, brain specific microRNA orthogroups (see Materials and Methods), results highlighted several neuronal (for example: “regulation of neuron projection development”, GO:0010975; “forebrain generation of neurons”, GO:0021872) behavioural(including “locomotory behaviour”, GO:0007626; “aggressive behaviour”, GO:0002118) and immune related processes (for instance, “negative regulation of innate immune response”, GO:0045824). Moreover, we find that the vast majority of these brain restricted orthogroups (32 out of the 55) have a species specific miRNA seed sequence (nt 2-8), potentially leading to novel regulatory interactions restricted to a particular lineage. Thus, our results suggest that the emergence of novel, tissue restricted microRNAs might play an important role in the lineage specific evolution of neuronal regulation, especially through the acquisition of novel seeds and associated targets.

### The co-evolution of microRNAs and their UTR target sites

Homology analyses and Dollo parsimony provided an overview of the evolutionary patterns of microRNA families in our species. Next, we asked the question whether we can detect signatures of selection acting on the predicted target repertoires. As a result of the selective constraints acting on microRNA target sites, we would expect these loci to show increased conservation compared to the surrounding 3’UTR regions. We compared *phastcons* scores of the targets associated with species specific and conserved seed families (defined as groups of microRNA loci sharing exactly the same miRNA seed sequence), and observed an evident increase in scores corresponding to the targets of conserved families (fig. 7). Additionally, the 3’UTR of genes targeted by conserved families show substantially higher conservation across all bins, as compared to those targeted by species specific seeds. These results suggest increased levels of purifying selection at the binding sites of conserved seed families.

**Fig 7.**
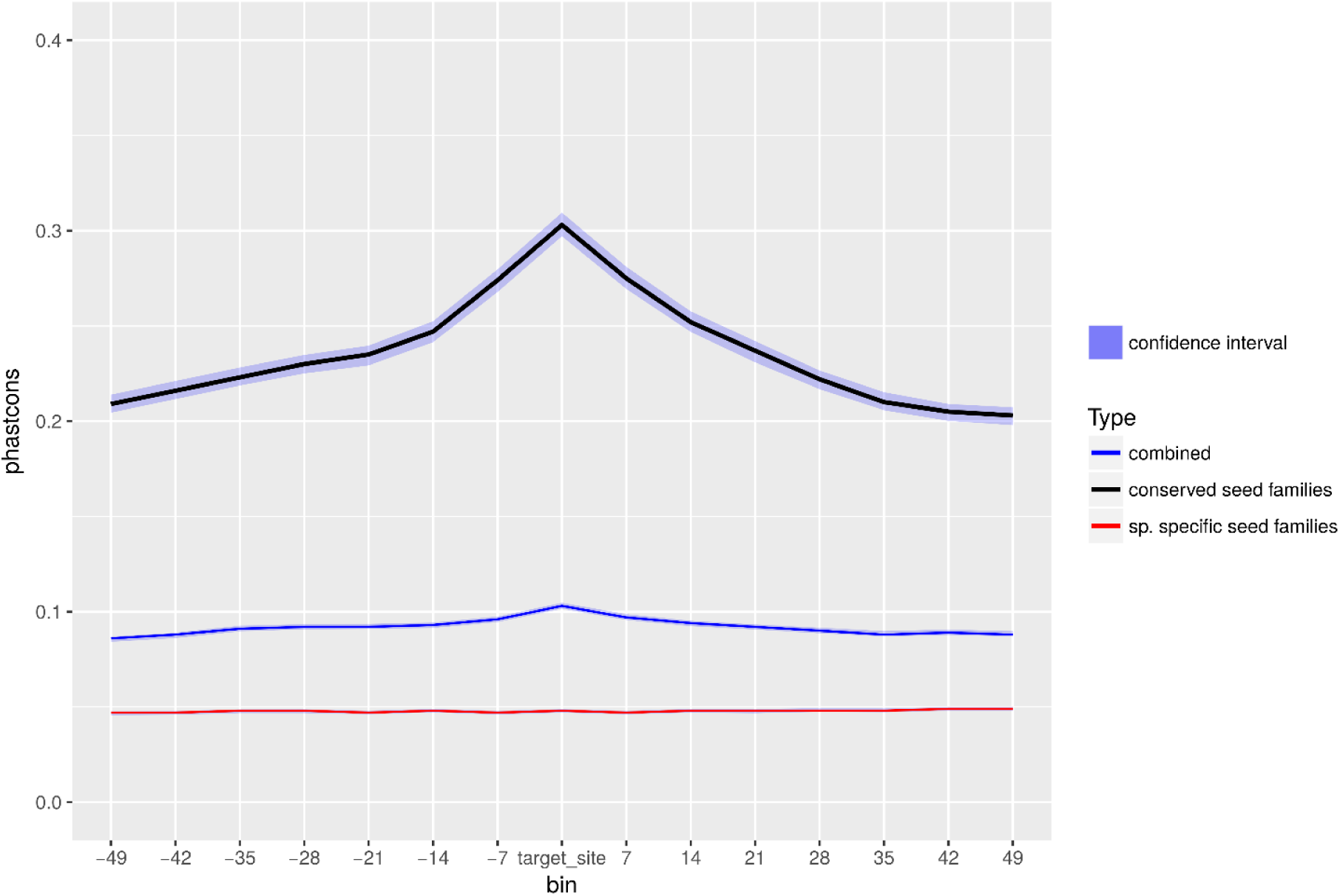
Median and confidence interval of 20-way Phastcons scores, calculated across 7nt bins, centred around the predicted targets of conserved (black line), species specific (red) and combined (blue) seed families.

We next asked the question whether 3’UTR target site conservation reflects the loss of a microRNA orthogroup (or seed family) during evolution. Based on homology analyses and Dollo parsimony inference, we first identified all microRNA orthogroups which appear to be lost in a terminal branch of our phylogenetic tree (Table 4) to test the hypothesis that the loss of a miRNA might be lead to relaxed selective constraints on the associated target sites.

We tested our hypothesis of relaxation of selection by calculating pairwise target sequence similarity between the rabbit (chosen as an outgroup) and all other species.

**Table 4.** Summary of the number of lost orthogroups (also absent in the miRBase annotation) and seed families.

|  | **Total**<br>**orthogroup**<br>**losses** | **Total**<br>**mature seed**<br>**losses** |
| --- | --- | --- |
| **Cow** | 9 | 13 |
| **Dog** | 2 | 1 |
| **Horse** | 4 | 7 |
| **Pig** | 3 | 4 |
| **Rabbit** | 18 | 15 |

However, we could not find any example where the species missing the microRNA family has a significantly lower conservation level in all comparisons (figg. S8-S13). The observed lack of evidence for differential target site conservation between the species retaining and losing the microRNA orthogroup could be explained by: 1) a very recent microRNA loss; 2) the seed sequence being shared with one or more conserved orthogroups leading to continued purifying selection; 3) the orthogroup loss having a very weak or null effect on target site conservation; 4) the presence of false positive target predictions in our data.

Seed sharing appears to be widespread in our set of species, as clearly shown in (fig.8). However, we also observe a considerable number of species specific seeds in the cow and the dog. Among the significantly enriched GO:term accessions (adjusted p-value<0.05) for dog specific seed families (fig.9) we find: “extracellular structure organisation” (GO:0043062), “positive regulation of axon extension” (GO:0045773), “positive regulation of neuroblast proliferation” (GO:0002052), and “behaviour” (GO:0007610). The significantly enriched GO:term accessions for cow specific seed families include “lactation” (GO:0007595) and “mRNA 3’ end processing” (GO:0031124), while terms found significant only for the dog include “forebrain development” (GO:0030900) and “DNA repair” (GO:0006281). Among the accessions enriched in both the cow and dog specific seed targets we find “behaviour” (GO:0007610), “behavioural fear response” (GO:0001662) and several immune-related processes: “T-cell activation” (GO:0042110), “T cell receptor V(D)J recombination” (GO:0033153), “negative regulation of innate immune response” (GO:0045824) and additional related accessions.

**Fig 8.**
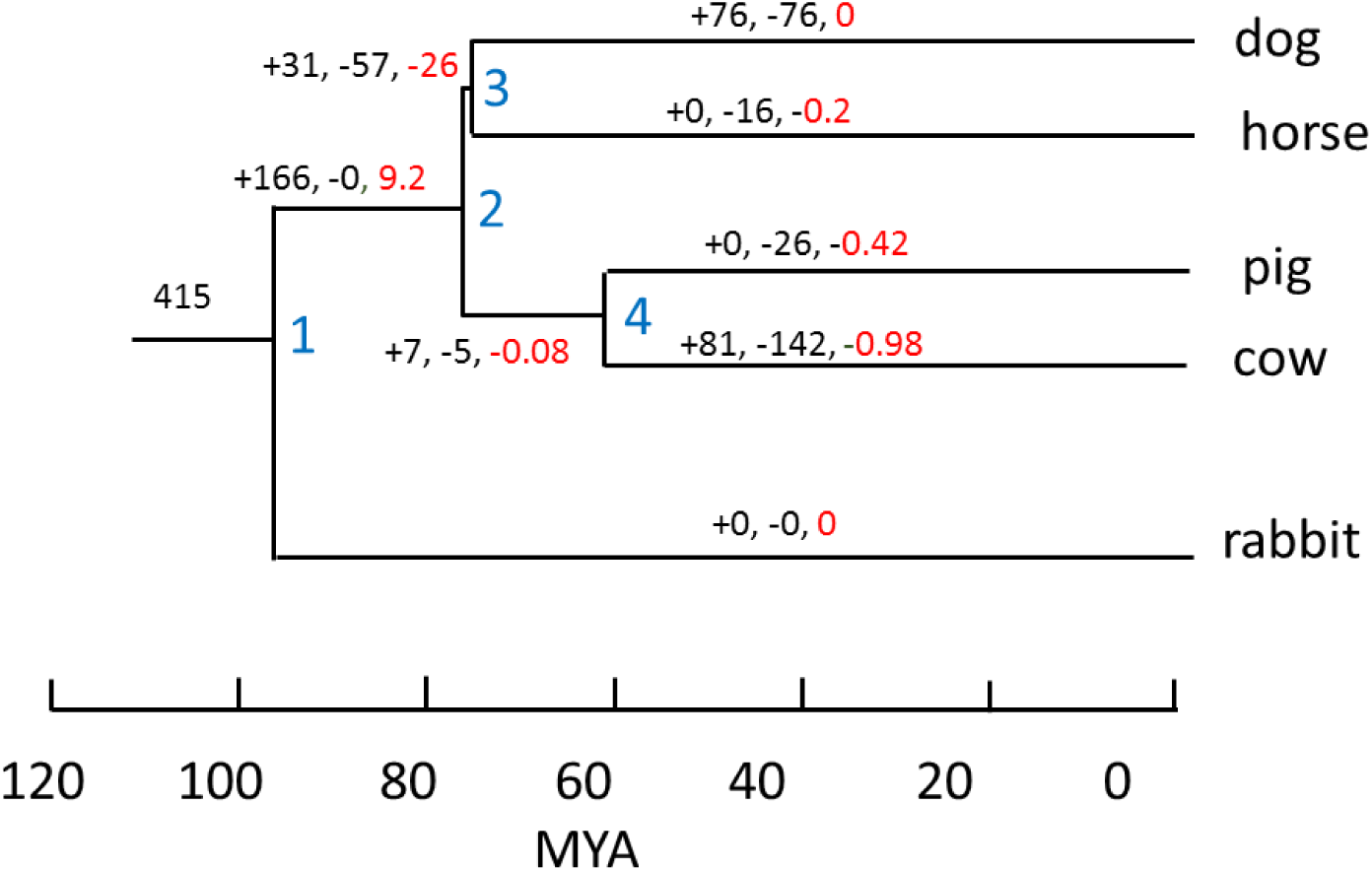
Gain and loss of miRNA seed sequences across the phylogeny.

**Fig.9.**
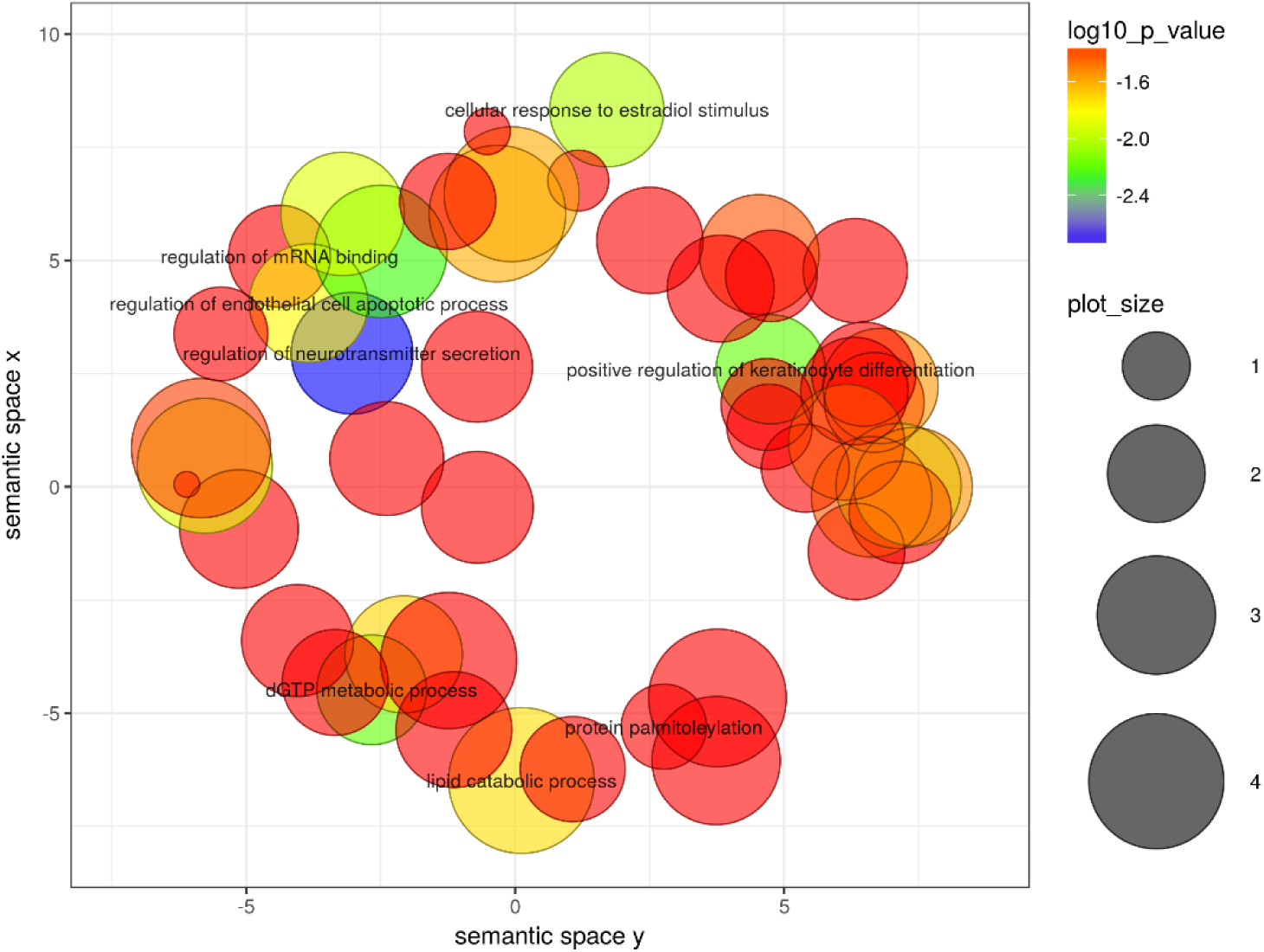
GO term results for dog specific seed family GGACCGA (orthogroup 490) as summarised by *Revigo* (Supek et al. 2011)

Next, we searched for domestication genes described in the literature (Boyko et al. 2010;) (Axelsson et al. 2013;) Freedman et al. 2016) in the target genes corresponding to significant GO accessions. 12 genes located in the candidate domestication regions identified by (Axelsson et al. 2013) are found among the targets of dog specific seed families. This set includes genes associated with behavioural (POLR1E), immunological (TLX3) and body weight (TNKS2) phenotypes in mouse (http://www.informatics.jax.org/).

When we considered the set of genes lying in the top 100 genomic regions under selection in the dog identified by *(Freedman et al. 2016)*, we found 26 genes belonging to one or more significant GO:term accessions for dog specific seed families. Once again, we observe genes associated with behaviour and body weight phenotypes (for instance GLRA1 and HTR2B). Despite a high number of predicted targets in the dog (3052, 91% of our target gene set), the overlap is significantly higher than expected (hypergeometric test, 1.27 fold enrichment, p=0.005)

We also looked for positively selected genes (Park et al. 2015);(Xu et al. 2015;)(Braud et al. 2017) in the set of targets of cow specific seed families belonging to a significant GO accession. We found 24 positively selected targets among the 35 common between our set of 1:1 orthologues and the gene list provided by (Park et al. 2015). (1.33 fold under-enrichment, p=10e^−4^). We observe genes associated with immune system, cardiovascular and muscular phenotypes, including TNFRSF8, CREBBP and MTMR14. Only 3 targets were found in the gene set from Xu *et al.* (Xu et al. 2015): WIF1 (Increased osteosarcoma incidence), KIT (coat/hair pigmentation) and LRIG3 (Abnormal craniofacial morphology) (2.43 fold under-enrichment, p=2x10e^−4^). Additionally, we considered the dataset from Braud *et al.* (Braud et al. 2017), representing genes with high miRNA binding sites divergence between *B.taurus* and *B. primigenius*. Among the top 200 scoring genes, we identified 43 targets of cow specific seed families (1.31 fold under-enrichment, p<10e^−4^). Even in this case, we observe many genes involved in immunity (for example CIITA, RHOH), brain morphology/behaviour/body size (ASPA), and neurological (FOXI1) phenotypes. Interestingly, FOXI1 (nervous system phenotype) appears as a positively selected (Freedman et al. 2016); (Braud et al. 2017) target of both dog and cow specific seed families.

Altogether, we clearly highlight a trend among positively selected microRNA targets towards a few biological processes and phenotypic traits. Thus, our results suggest that species specific seed families might have played a role in domestication, by modulating the expression of genes under artificial positive selection.

## Discussion

Our study provides for the first time a comparative analysis of microRNA and target evolution across five domestic mammals. Our data allows us to generate an improved microRNA annotation across five mammalian genomes and four different tissues. We take advantage of this dataset to investigate the potential contribution of microRNAs to the domestication process, the molecular mechanisms underlying the birth of novel microRNAs, and their co-evolution with 3’UTR target sites.

Our data highlights the importance of intronic sequences (366 orthogroups), duplication events (135 orthogroups) and repetitive elements (37 orthogroups) for the emergence of new microRNA genes. We provide evidence for high levels of microRNA turnover across the phylogeny, and a general increase in net gain rates along terminal branches. With the only exception of the horse, these rates are comparable to previous studies on mammalian microRNA evolution (Meunier et al. 2013).

Novel microRNAs tend to be more tissue specific compared to the miRBase set, with the most striking difference being represented by the brain tissues. Indeed, we observe that 20% of novel orthogroups are restricted to the brain tissues, and the associated target repertoires are enriched for behaviour, neuron activity and differentiation processes. Similar results are observed for the targets of repeat derived, dog specific microRNAs.

Branch specific losses of a seed or an orthogroup appear to be rare events, and do not result in a detectable inter species difference in sequence conservation. This holds true even when we consider the evolution of microRNA orthogroups.Additionally, we find examples of microRNA loss compensation through the retention of loci with exactly the same seed sequence. The lack of evidence for differential target site conservation between the species retaining and losing a microRNA family can be explained in many ways: a very recent microRNA loss; the lack of orthogroup seed specificity (when considering orthogroups rather than seed families); the presence of false positives in our target predictions, or the absence of any effect on target site conservation.

An additional factor to consider is represented by the demographic history of our five species. Population bottlenecks associated with the domestication process can have a severe impact on genetic diversity, affect the strength of natural selection, as well as determine an increased accumulation of mildly deleterious mutations (Carneiro et al. 2011;)(Schubert et al. 2014;)(Alves et al. 2015;)(Scheu et al. 2015;)(Marsden et al. 2016).

While a vast proportion of miRNA sequences are conserved, our dataset provides evidence for the emergence of species specific seed sequences in the cow and the dog. GO term analyses of the associated targets highlight the enrichment for neurological, behavioural and immune related processes. Moreover, these target repertoires include several genes which have been described as positively selected during the domestication process. Thus, our results suggest an involvement of specific seed families in the domestication process by modulating the expression of positively selected genes, thus contributing to morphological and behavioural variation, neurological processes and immune response.

Compared to protein-coding genes, microRNAs represent a relatively simple source of innovation, as they can rapidly evolve from already transcribed genomic regions. Short sequence similarity is sufficient for mRNA targeting and down-regulation, while the turnover of miRNA seed sequences and target sites provides the organism with a wide space of possible regulatory changes. While a significant fraction of microRNAs appear to be conserved over long evolutionary times, our data confirms previous observations of high evolutionary turnover in animals, with many orthogroups appearing to be lineage specific (Berezikov 2011); (Meunier et al. 2013;) (Flynt 2017). The gain of novel microRNAs might result in the acquisition of novel regulatory pathways, and spatio-temporal changes in protein coding gene expression. It represents an additional layer of regulatory complexity which we are still trying to fully uncover. Further research is needed to better clarify the extent of the contribution of microRNAs to lineage specific adaptations and phenotypic diversity.

## Materials and Methods

### Small RNA library sequencing

Tissue samples were obtained commercially from *Zyagen*. Heart, kidney and testis were obtained for all five species. As for the brain tissues, we obtained four different brain regions (cortex, cerebellum, hypothalamus) for the cow, the dog and the pig, and whole brain for the horse and the rabbit. Small RNA libraries were prepared using the *TruSeq Small RNA Library Prep Kits*. six-base indexes distinguish samples and allow multiplexed sequencing and analysis using unique indexes ( (Set A: indexes 1 - 12 (RS-200-0012), Set B: indexes 13-24(RS-200-0024), SetC: indexes 25-36 (RS - 200 - 0036), (TruSeq Small RNA Library Prep Kit Reference Guide, Part 15004197 Rev.G).

The TruSeq Small RNA Library Prep Kit protocol was followed using an input of 1µg of total RNA.

Quantification of total RNA was done using the Qubit RNA HS Assay kit*(ThermoFisher* Q32852).

RNA quality was established using the *Bioanalyzer RNA Nano kit* (Agilent Technologies 5067-1511).

An RNA Integrity Number (RIN) value ≥8 was required for the RNA to pass the QC step.

This protocol generates small RNA libraries directly from RNA by ligating adapters to each end of the RNA molecule. Reverse transcription is used to create cDNA, and PCR amplification of the cDNA (14 cycles of PCR in the standard protocol) is used to generate libraries. Library purification combines the use of *BluePippin* cassettes *((Sage Science Pippin Prep 3%* Cassettes Dye-Free(CDF3010), set to collection mode range 125 - 160bp)) to extract the library molecules with a concentration step *(Qiagen MinElute PCR Purification* (cat. no. 28004)) to produce libraries ready for sequencing. Library concentration and size are established using *HS DNA Qubit* and HS DNA *Bioanalyser*.

All libraries were pooled together and sequenced on 2 lanes of an Illumina *HiSeq* machine. With the only exception of dog cortex, all libraries were later sequenced on a second run of the Illumina *HiSeq*, using the same pooling strategy. This run, however, was loaded with 10% *phix* to increase sequence diversity, which lead to an improved read quality.

In the case of dog cortex, two additional libraries were also constructed (as part of the development work establishing the protocol) with a different mix of conditions: 11 PCR cycles + PAGE, and 14 PCR cycles + PAGE. All 3 available dog cortex libraries (11 PCR cycles % PAGE, 14 PCR cycles % PAGE and 14 PCR cycles % Pippin) were sequenced on an Illumina *MiSeq* machine, and resulting sequencing data was included in the study.

### Annotation of conserved and novel microRNAs

For each organism, we ran *miRCat (Stocks et al. 2012)* and *miRDeep2 (Friedlander et al. 2012)* on the corresponding combined set of small RNA libraries, thus generating two independent sets of putative miRNA loci. Genomic coordinates of miRCat and MiRDeep2 predictions were then merged using *Bedtools merge* in order to generate a non-overlapping set of loci.

We then aligned small RNA reads of each library to our predicted hairpins. We looked for evidence of 3p-miRNA and 5p-miRNA expression in the alignments, and generated hairpin secondary structures as well using the *Vienna-RNA* package. Based on the consistency of both the alignments and predicted secondary structure, a set of high confidence miRNA loci was derived.

Initially, all loci covered by less than 10 reads were discarded. However, some of these genes were rescued at a later stage, when we generated BLAST alignments of our final set of microRNAs against the 5 genomes (see *Homology and synteny analyses)*. Specifically, when the low-coverage prediction showed both evidence of Dicer and Drosha processing and sequence homology (as identified by the BLAST analysis) with a microRNA gene present in our annotation, the gene was rescued and added to the final dataset.

The identification of novel microRNA loci was performed as follows. First, we aligned the complete set of miRBase hairpin sequences to the organism’s genome sequence, using the command-line version of *BLASTN* (e-value ≤ 10-^6^)(Altschul et al. 1990). We then mapped all mature miRBase sequences to these putative pre-miRNA hairpins, and selected those for which at least one alignment with no more than one mismatch was observed. The subset of novel microRNA genes was then identified by removing all predicted miRNA loci overlapping at least one miRBase genomic hit. Tissue specific expression plots were generated using *Rstudio* (https://www.rstudio.com/).

### Homology and synteny analyses

The combined set of annotated miRNA loci was aligned, using BLASTN (Altschul et al. 1990), against the latest genome assemblies for our five species, as well as human and mouse. We selected BLAST hits with an e-value ≤10-^6^ and an alignment length of at least 40 nt. We then identified the closest protein coding gene upstream and downstream of the selected hits, as well as the gene containing the hit for all intragenic hits. Genes surrounding or containing the query (microRNA) and the subject (BLAST hit) sequences were compared, looking for the presence of at least one homologous pair of genes with conserved synteny structure (i.e. same upstream or downstream gene, both on the same or on the opposite strand with respect to the microRNA/BLAST hit). Any BLAST hit supported by synteny conservation of at least one protein coding gene was then flagged as an orthologous locus. Predicted microRNA loci across all five species were grouped into orthogroups, using *CDhit* (Fu et al. 2012) with 80% minimum identity.

To characterise the most likely patterns of gain and loss of microRNA clusters across the phylogeny, we ran dollop from the package phylip-3.696 (http://evolution.genetics.washington.edu/phylip) on the 732 microRNA orthogroups.

### Identification of repeat-derived microRNAs

We used BLASTN to align the hairpin sequences of our annotated microRNAs against all sequences in the *Repbase* database (http://www.girinst.org/repbase). We then selected all repetitive elements which had been returned by our BLASTN search, and generated 1000 shuffled sequences for each of these elements. All microRNAs were subsequently aligned (using BLASTN) to all of these shuffled sequences. Finally, we selected all original BLASTN results having a minimum alignment length of 30 nt, as well as a bit score higher than the maximum value (42.8) observed in the alignments against the shuffled repetitive sequences.

### Generation of 3’UTR multiple alignments

The latest genome annotation for our 5 species was downloaded from the Ensembl website (www.ensembl.org). Genes representing one to one orthologs across all 5 species plus human and mouse were selected. The corresponding UTRs were defined as the region starting from the end position of the last annotated exon, plus one, and ending 5kb downstream. Any overlap with downstream coding sequences, as determined using *bedtools subtract* (Quinlan 2014), was then removed by trimming the 5kb window up to the starting position of the first overlapping coding sequence. Resulting UTR sequences were then filtered for a minimum length of 500 bp. We used *mafft* (Katoh and Standley 2013) to generate gene specific multiple alignments across our 5 species, as well as human and mouse. The final alignments were obtained by extracting the alignment region corresponding to the human or the mouse homologous sequence, depending on which one was the longest sequence. The final dataset used for our analyses includes 3355 one to one orthologues across 7 species.

### Target prediction and Gene Ontology analyses

Target sites were identified in silico using *TargetScan7* (Agarwal et al. 2015), limiting the set of targeted genes to the 1:1 orthologues across all five species. The sequences of all predicted 7mer and 8mer miRNA-target interactions were then extracted. For the analyses of branch specific microRNA and seed losses scenarios, we predicted target sites across all our five species. We then selected all 7mer and 8mer sites which were independently predicted in the outgroup (rabbit) and at least one additional species, with an associated *context++* score (calculated without *Pct* contribution)(Agarwal et al. 2015) smaller or equal to −0.1. Target sites sequences plus 2 nucleotides upstream and downstream were aligned using *mafft* (Katoh and Standley 2013). Multiple alignments were then used to calculate the pairwise sequence similarity across target sites between pairs of species. In order to consider all target sites at once, sequence similarity was calculated on the 6mer sequence complementary to positions 2-7 of the miRNA.

We adapted the above described pipeline for the calculation of *Phastcons* scores across 7nt bins in the 3’UTR regions. For conserved seed families, we selected targets sites predicted in at least 2 species, and with an associated *context++* score smaller or equal to −0.1. For the species specific seeds, we onlyusedthe *context++* score as a filtering criterion. Sequences corresponding to the target site ± 49nt were identified, and the genomic coordinates of the homologous human counterparts inferred from multiple alignments. We then used a combination of *UCSCtools (Kent et al. 2010)* and in-house perl scripts to calculate single nucleotide *Phastcons* scores across bins, for each target site of every seed family.

Gene Ontology analyses were performed in *Rstudio* using the package *topGO* (https://bioconductor.org/packages/release/bioc/vignettes/topGO/). Significantly enriched GO terms were independently identified for the targets of each seed family, using the *elim* algorithm coupled with Fisher exact test. The gene background was defined as the complete set of 1:1 orthologues used for all target sites analyses. For the GO analysis of brain specific novel microRNAs, this background was restricted to the target genes with evidence of expression of the human homologue in brain tissues (https://www.proteinatlas.org/about/download). The raw p-values obtained were not corrected, as the *elim* method already includes an adjustment equivalent to a Bonferroni correction (Alexa et al. 2006). Results were filtered for p≤0.05.

## Acknowledgements

SM, WH and FDP were supported by the BBSRC, Institute Strategic Programme Grant [BB/J004669/1]; LPD and SM were supported by the Functional Annotation of Animal Genomes [BB/M01844X/1]; WH and FDP are supported by the BBSRC Core Strategic Programme Grant [BB/P016774/1].

Next-generation sequencing and library construction was delivered via the BBSRC National Capability in Genomics and Single Cell (BB/CCG1720/1) at Earlham Institute by members of the Genomics Pipelines Group.

